# Comprehensive mRNA annotation in trypanosomatid parasites

**DOI:** 10.64898/2026.02.24.707742

**Authors:** Ulrich Dobramysl, Richard J Wheeler

## Abstract

Trypanosomatid parasites, including human infective *Leishmania* and *Trypanosoma* species, have an unusual genome organisation and transcription. They are unicellular eukaryotes, but unlike most eukaryotes, which have individual promoters per gene, most protein coding genes are co-transcribed in long gene arrays. This nascent transcript is processed into individual mRNAs by trans-splicing and polyadenylation. Accurate analysis of transcription, transcript processing and transcript abundance requires accurate genome annotation of spliced leader acceptor sites, polyadenylation sites and the resulting 5′ and 3′ mRNA untranslated regions. Here, we describe tools for annotating these features from short read RNA sequencing data and for measuring the usage of spliced leader acceptor and polyadenylation sites. These are practical, scalable software packages, and we use them to annotate UTRs across all available trypanosomatid genomes.

## Introduction

The trypanosomatids are a group of professional unicellular parasites which includes the human *Leishmania, Trypanosoma cruzi* and *Trypanosoma brucei* pathogens, numerous *Trypanosoma* and *Leishmania* animal pathogens, along with a wider diversity of plant and monoxenous insect parasites. Together they are responsible for an enormous health and economic burden to tropical and sub-tropical regions globally.

There is ever-growing genomic data for this family. Most genomes are hosted on the community-run TriTrypDB database, part of the wider VeuPathDB project (Amos et al., 2022). As of version 68, TriTrypDB host 88 genomes across 38 species, recording the moderately large and complex genomes which encode the capacity to be successful pathogens. However only 3 genomes have a large number of annotated 5′ and 3′ mRNA untranslated regions (UTRs): *Trypanosoma brucei* TREU927, EATRO1125 and *Leishmania major* Friedlin 2021. Of these, only *T. brucei* TREU927 is close to complete.

Improved UTR annotation would greatly benefit our understanding of gene expression control in these parasites. In trypanosomatids, almost all protein-coding genes are co-transcribed in long arrays containing many tens to hundreds genes (Berriman et al., 2005; Daniels et al., 2010), and subsequently processed into mature mRNAs by trans-splicing and polyadenylation (Ullu et al., 1993), with only rare instances of cis-splicing (Mair et al., 2000). Trans-splicing adds the spliced leader (SL) to the 5′ end at the spliced leader acceptor site (SLAS), and polyadenylation adds the poly-A (PA) tail to the 3′ end at the polyadenylation site (PAS). This form of transcription limits the capacity for transcriptional control, with mRNA stability (particularly conferred by 3′ UTR sequences) canonically thought to dominate (Pays, 2005). Lack of consistently annotated UTRs (both on TriTrypDB and elsewhere) inhibits research into this area, from hampering bioinformatic analysis of conserved regulatory UTR elements to restricting RNA-seq analysis of transcript abundance to use the CDS rather than full transcript.

PASs and splicing are readily detectable from RNA-seq data (Wang et al., 2009). In the *T. brucei* TREU927 genome project (Berriman et al., 2005), the first complete trypanosomatid genome, UTRs were later mapped using a combination of PA-enriched RNA-seq, ribosome footprinting and SL-enriched short read RNA-seq (Jensen et al., 2014; Nilsson et al., 2010; Parsons et al., 2015; Siegel et al., 2010). Much more recently, UTRs were mapped in *L. major* Friedlin 2021 from short read RNA-seq data by identifying reads containing SL or PA sequences (Camacho et al., 2021) and *de novo* transcript assembly from pooled single cell RNA-seq data was used to map the *T. brucei* EATRO1125 UTRs (Naguleswaran et al., 2021). However, UTR annotation information does not always reach the genome databases. As examples, neither the older short read RNA-seq *L. major* (Dillon et al., 2015) and *L. mexicana* (Beneke et al., 2022; Fiebig et al., 2015) studies, nor the recent long and short read RNA-seq *L. infantum* (Camacho et al., 2023) study are represented within TriTrypDB.

The consensus from these studies is that SLASs and PASs can be efficiently found from short read RNA-seq data: reads which end with either the spliced leader sequence or the PA tail are identified, the SL/PA trimmed, and the remainder of the read mapped to the genome to identify the SLAS and PAS sites. We are aware of two tools which are based on this methodology, SLaPmapper (Fiebig et al., 2014) and UTRme (Radío et al., 2018), however neither has led to trypanosomatid UTR data becoming widely available.

It is possible to predict PASs and SLASs from the genomic sequence alone as they are located near a polypyrimidine tract with splicing occurring at a characteristic dinucleotide (Benz et al., 2005). Tools using this RNA-seq-independent computational methodology have been developed (Gopal et al., 2005; Kelly et al.; Smith et al., 2008), however, as they were developed prior to much contemporary training data being available their predictions lack accuracy.

SLs and PA are natural parts of all mRNAs, hence SLAS/PAS information will be present regardless of the specific RNA-seq sample preparation methodology. Trypanosomatid-specific library preparation strategies which enrich in SL containing RNAs (Nilsson et al., 2010) may bring advantages, while there are also RNA-seq sample preparation-specific artefacts (Verwilt et al., 2023). Nonetheless, the sheer quantity of RNA-seq data now available across different species, despite being captured without SLAS/PAS mapping having been a design consideration, allows a data-based annotation strategy rather than relying on sequence-based prediction.

Therefore, we developed `slapquant`, a tool which maps SLASs, PASs and UTRs with minimal user input from standard short read RNA-seq data. Its primary usage mode is to map the UTRs for genomes with annotated CDSs using RNA-seq data, improving genome annotation and enabling downstream analyses of evolution of SLAS/PAS sites, UTR regulatory elements, etc. We apply this tool to annotate UTRs in 47 genomes that have CDSs annotated and are available on TriTrypDB. This tool and dataset open numerous avenues for future study, such as: a direct analysis of SLAS/PAS sites, for example considering life cycle stage-specific SLAS/PAS (similar to previous *Leishmania* promastigote and amastigote comparison (Dillon et al., 2015; Fiebig et al., 2014)); identifying when SLAS/PAS sites fail to be used (as we previously used to identify ESB1 as a transcriptional regulator (López-Escobar et al., 2022)); and any downstream study of mRNAs, from improving transcript quantitation where knowing the full-length mRNA sequence is important, to any study involving analysing UTR sequence involvement in regulating transcript abundance and translation efficiency.

## Results

We implemented SLAS and PAS detection, assignment to CDSs and UTR annotation in Python, requiring either BWA MEM (Li, 2013) or BWA MEM2 (Vasimuddin et al., 2019) for alignment and using AWK for fast analysis of the output, broken down into tools which can be used sequentially:

`slapquant`, taking FASTQ file(s) of short read (eg. Illumina) sequencing reads and the reference genome fasta file, and outputting a genome functional format (GFF) file containing the locations of identified PASs and SLASs and their usage incidence.

slapassign, taking the GFF file from slapquant and the reference genome GFF with protein coding genes with CDSs annotated, and outputting a GFF file with PASs and SLASs assigned to consistent protein coding genes.

slaputrs, taking the GFF file from slapquant and adding 3′ and 5′ UTRs to protein coding genes, optionally correcting the CDS of the gene based on SLAS / PAS position.

Additionally, we developed `slapspan`, which takes a GFF file from `slapassign`, or `slaputrs`, the reference genome and FASTQ file(s) of sequencing reads and returns statistics of how many reads span SLASs or PASs. These spanning reads likely originate from nascent transcripts and thus provide essential information about unprocessed RNA.

### Trial T. brucei and L. mexicana datasets

We used two trial species for optimisation. *Trypanosoma brucei* TREU927 using publicly available RNA-seq data not captured with the purpose of SLAS and PAS mapping in mind and *Leishmania mexicana* MNYC/BZ/62/M379 using sequencing data captured with the purpose of SLAS and PAS mapping in mind:

For *T. brucei*, we used all TREU929 procyclic form data from our recent study (López-Escobar et al., 2022) and the reference genome (Berriman et al., 2005). For *L. mexicana,* we used the promastigote and axenic amastigote data previously used to identify SLAPs, PAS, and map the UTRs (Fiebig et al., 2015) and our strain-specific genome (Beneke et al., 2022). All are 100 bp paired end datasets with short insert size. For *T. brucei,* libraries were random primer PCR selected, and for *L. mexicana,* a mixture of random and poly-T selected.

### Optimisation and performance of spliced leader acceptor and polyadenylation site detection using slapquant

`slapquant` carries out detection of SLAS and PAS sites. Previous approaches took short sequencing reads and filtered them to identify those which end in a 5′ SL, 3′ SL reverse complement, 3′ PA or 5′ PA reverse complement (poly-T), trimmed these read to remove the SL/PA sequence, and aligned the remainder of the read to the reference genome. The trimmed end of the read is then taken as the site of a SLAS/PAS (Fiebig et al., 2014; Radío et al., 2018). Here we instead align all full-length reads to the reference genome, identify which reads have a 5′ or 3′ which fail to align while the remainder of the read does (clipped read alignments). For each clipped alignment, we ask whether the sequence immediately next to the clipping matches a 5′ SL, 3′ SL reverse complement, 3′ PA or 5′ PA reverse complement, and, if it does, then take that as a SLAS/PAS (Figure 1). This avoids a set of potential false positives: Consider, for example, a genome-encoded PA sequence which is transcribed as part of an mRNA. Reads which happen to have their 3′ end in that PA would be falsely detected as PA-containing by the read filtering strategy, leading to incorrect mapping of a PAS at the start of the PA sequence. Previous tools use post-alignment checks of the genome-sequence for containing PA (Fiebig et al., 2014) or more complex similarity and A-richness scoring (Radío et al., 2018) to correct this error. However, these will be correctly rejected by our clipped alignment strategy.

**Figure 1.**
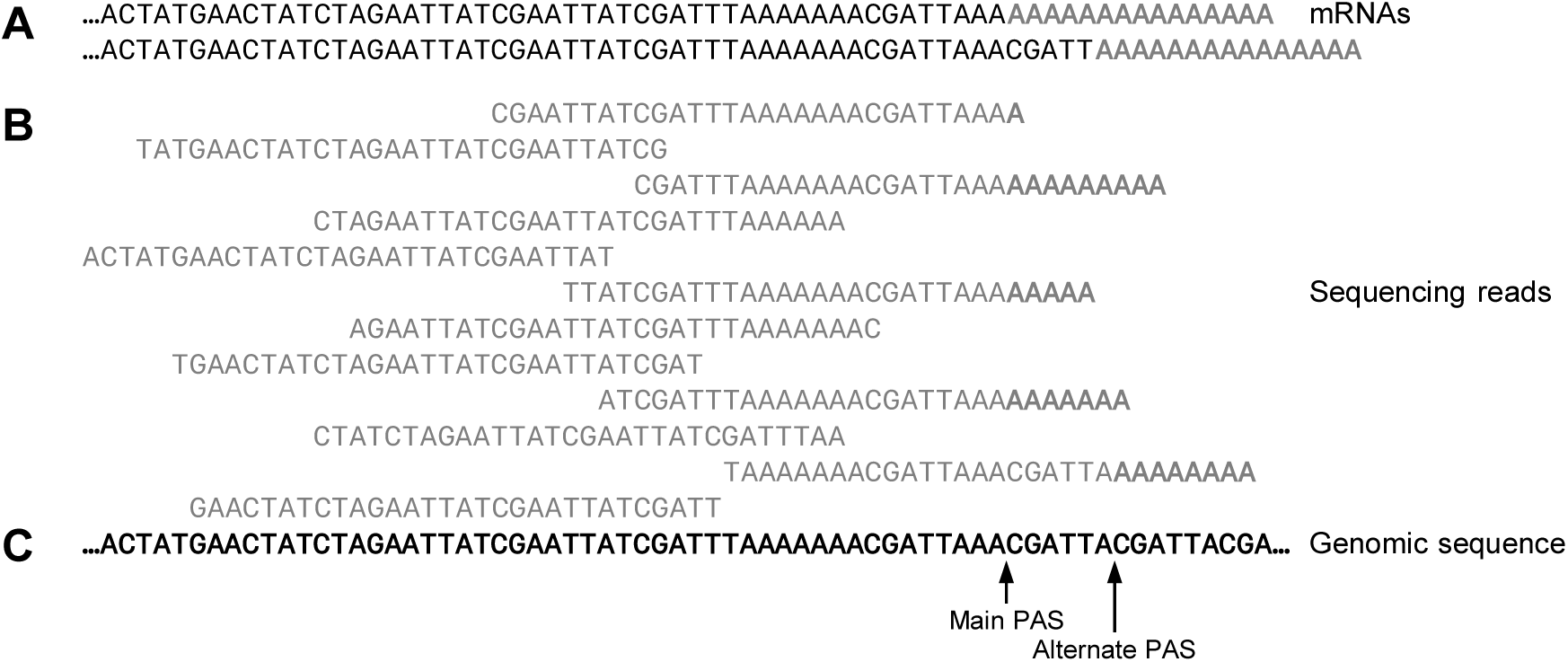
Pileup strategy for detection of SLASs and PASs, illustrated using a PAS Illustrative example, using 30 bp reads, of sequencing reads from two mRNAs aligned to the genomic sequence to detect a main and an alternate polyadenylation site (PAS). **A.** The mRNAs, with short illustrative poly-A tails. **B.** Illustrative sequencing reads, aligned to the genomic sequence. Mismatches are marked in bold. When matching poly-A the point at which mismatches start is taken as the PAS. (Note that the illustrative reads and poly-A mismatch regions are shown shorter than in reality for clarity). **C.** Genomic sequence, indicating where poly-A leads to mismatches at the end of the sequencing alignment (alignment clipping) indicating a polyadenylation site.

We implemented `slapquant` to handle sequencing read alignment and directly analyse the output from BWA MEM or BWA MEM2, to eliminate the need for storage or slow write and read of a large intermediate sequence alignment map (SAM) or binary alignment (BAM) file. Thus, the necessary input is the reference genome sequence in FASTA format, and the sequencing read file(s) in FASTQ or gzipped FASTQ format, and the output is a General Feature Format (GFF) file containing the locations of SLASs and PASs.

`slapquant` has two tuneable parameters for detection of PAS and SLAS sites, the minimum length of exact sequence match in the clipped sequence to be considered a PA or SL match. We asked how both total number of detected sites and total usage of those sites depended on match length. Following a drop from non-specific identification with very short (<5 bp) SL match length, there was a clear plateau of number and usage of candidate SLASs (Figure 2A). For PA match, there was a drop from non-specific identification with very short (<5 bp) PA match length, beyond which there was an approximately exponential decrease of number and usage of candidate PAS sites detected with length of PA match (Figure 2A). This suggests specific SL and PA matches for match lengths ≥6 bp. Consistent with previous observations, mean usage of SLASs was higher than PASs, indicating more uniform use of few SLASs and more diversity in selection of PASs.

**Figure 2.**
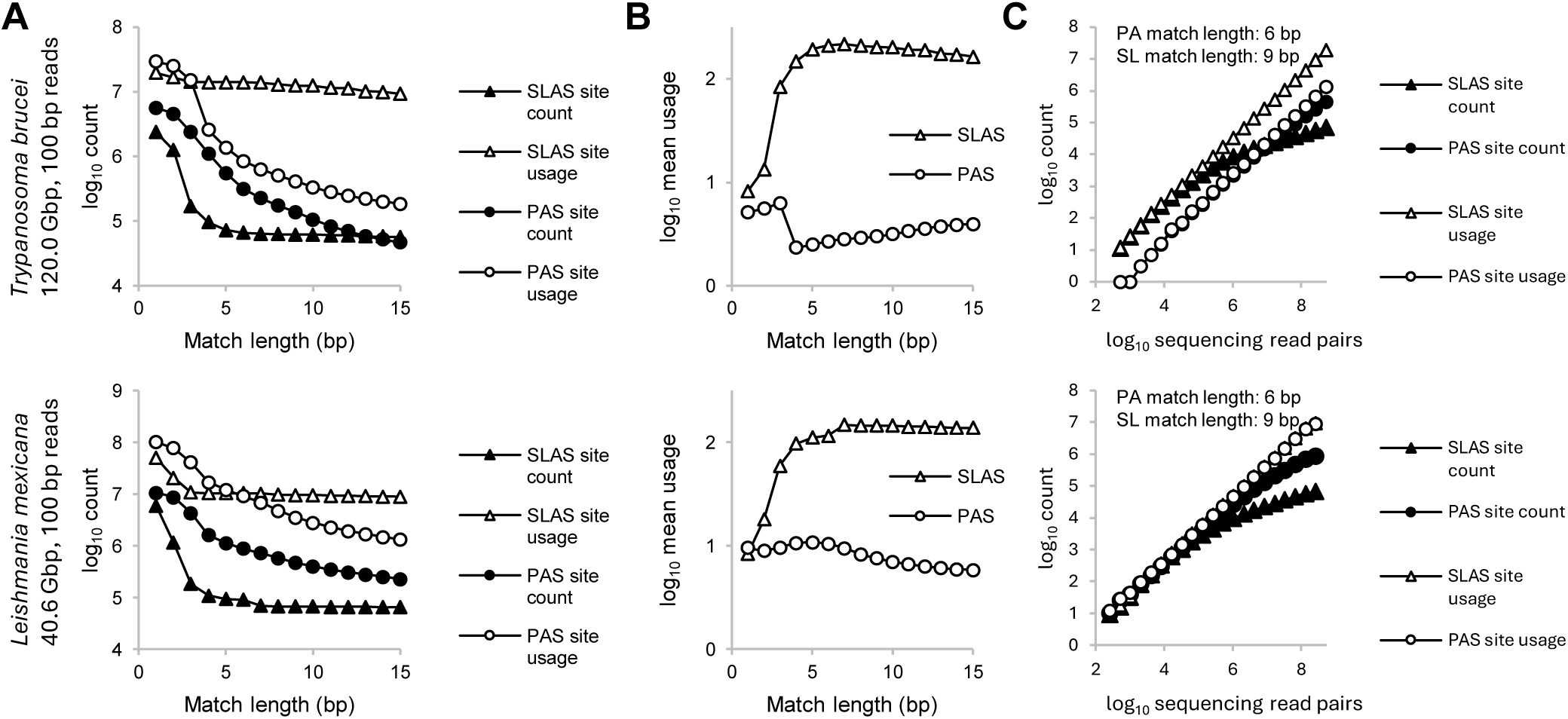
Benchmarking SLAS and PAS detection and usage rate **A.** The number of spliced leader acceptor sites (SLASs) and polyadenylation sites (PASs) detected, and the corresponding total SLAS and PAS site usages observed, for different minimum match lengths. Determined for (top) *T. brucei* and (bottom) *L. mexicana* using paired end 100 bp read sequencing data. **B.** The mean number of observed usages of each SLAS and PAS site for different minimum match lengths, corresponding to total site usage divided by number of sites for SLASs and PASs respectively in A. Determined for (top) *T. brucei* and (bottom) *L. mexicana.* **C.** Number of SLASs and PASs sites detected, and corresponding total SLAS and PAS site usages observed, for different numbers of sequencing reads. Detected using 6 bp and 9 bp match length for PAS and SLAS, using sequencing data randomly subsampled from that used for A and B.

These plots can be interrogated to determine potential sensitivity of the input dataset. The clear independence of number of detected SLAS sites and their usage with SL match length supports high accuracy SLAS detection, corresponding to the number of mRNA 5′ ends contributing to the dataset, ∼10,000,000 for both the *L. mexicana* and *T. brucei* datasets respectively. Assuming no 5′ or 3′ bias in the sequencing, equal usage of PASs corresponding to the same number of mRNA 3′ ends would be expected – although real datasets deviate from this ideal. For *L. mexicana*, a PA match length of 6 bp gave PAS site usage approximately equal to SLAS site usage, so PAS detection from ∼10,000,000 mRNA 3′ ends. For *T. brucei*, PAS site detection was underpowered, with PAS usage dropping below SLAS usage by a PA match length of only 4 bp, and ∼20× lower at our suggested minimum match length of 6 bp. Thus, PAS were detected from only ∼500,000 mRNA 3′ ends.

These quantitative evaluations of performance guided us to select a default PA match length of 6 bp and SL match length of 9 bp, as a balance between sensitivity and match number. However, both can also be set to custom values (using `slapquant --sl-length` and `--pa-length`, and our strategy here can be used to guide selection of these values.

### Spliced leader sequence identification using slapidentify

SLAS identification using `slapquant` requires an SL sequence. This can be provided as a previously determined sequence, or we provide built-in SL sequences for commonly analysed trypanosomatid genera, from our comprehensive analysis of different species below. We consistently observed CAGTTTCTGTACTTTATTG for *Leishmania* and CAGTTTCTGTACTATATTG for *Trypanosoma*. If no sequence or species is provided, `slapquant` attempts to identify the likely SL from the most common sequence at the end of reads, ignoring PA. This is also implemented as the standalone `slapidentify` tool. However, this approach is sensitive to incorrectly trimmed reads, which have sequencing adapters at the 5′ end, or non-SL sequences which are common due to the biology of the organism. For example, when searching for SL sequence lengths from 5 to 15 bp in our trial datasets (with no read trimming or other quality filtering) the *T. brucei* SL sequence is correctly identified (for all search lengths) but the *L. mexicana* SL sequence was not.

When an SL sequence is unknown, we recommend, at minimum, careful critical evaluation of automatic SL sequence identification in `slapquant` or the output from `slapidentify`. The preferred approach is using known SL sequences or to use dedicated SL identification tools (Calvelo et al., 2020; Wheeler, 2021a).

### Strategy for spliced leader and polyadenylation sites assignment to coding sequences using slapassign

SLASs and PASs define the 5′ and 3′ end of the genome-encoded portion of mRNAs, and, due to almost complete lack of introns, trypanosomatid genome annotation typically uses a CDS detection-first pipeline. Thus we implemented `slapassign` to assign SLASs/PASs to CDSs and `slaputrs` to annotate 5′ and 3′ UTRs using this information. This does, however, rely on accurate CDS annotation.

Previously described assignment of SLASs/PASs to CDSs simply assign all sites to the next downstream (SLAS) or upstream (PAS) CDS (Fiebig et al., 2014). However, this makes errors when unannotated genes or pseudogenes are transcribed and processed from this intergenic space.

Here, aware that there is a large range of mRNA abundances and thus a large range in the number of SLAS/PAS sites detected per CDS and their observed usage rate, we used an assignment strategy which adapts to the local SLAS/PAS site usage. First, we filter SLASs and PASs to only include those observed as used at least twice, as a simple means to eliminate rare biological or technical noise. Minimum SLAS and PAS usage can also be independently set by the user (`slapassign --min-slas-usage` and `--min-pas-usage`). Second, for each SLAS or PAS, we search downstream and upstream respectively for the next annotated CDS, and check for intervening sites: PASs between a candidate SLAS and its CDS, and SLASs between a candidate PAS and its CDS. If the usage of the candidate SLAS/PAS is greater than any intervening PAS/SLAS sites, then the candidate SLAS/PAS site is accepted and assigned to the CDS. Finally, the 5′ and 3′ UTRs are assigned from the usage weighted median of the CDS-assigned SLAS and PAS sites respectively (Figure 3).

**Figure 3.**
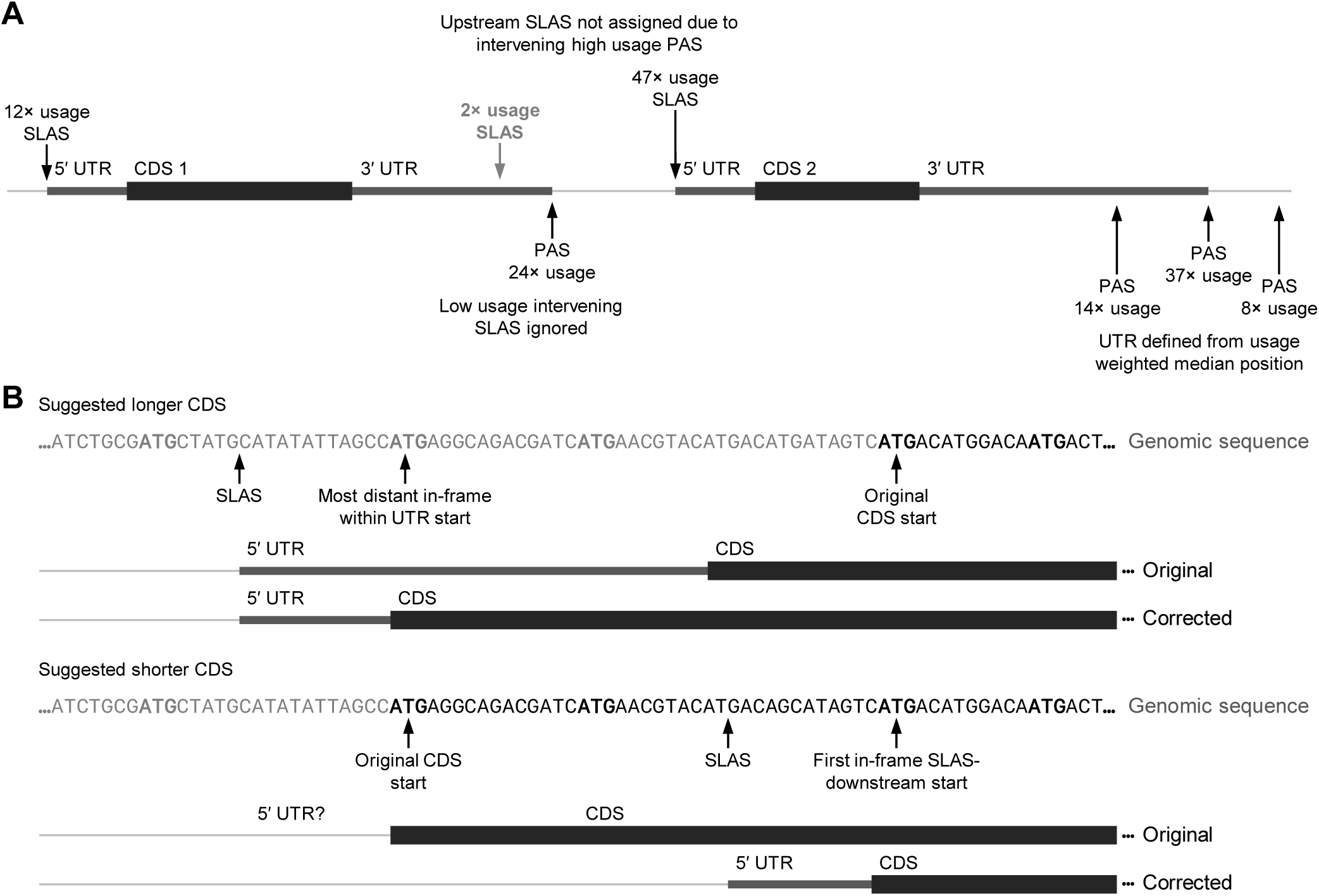
UTR definition from SLAS and PAS sites **A.** Simple illustrative example of UTR definition for two genes running right to left. When a single SLAS or PAS is assigned to the next downstream or upstream CDS, this defines the start or end of the 5′ or 3′ UTR respectively, for example CDS 1 5′ UTR. When multiple SLASs or PASs are assigned to a CDS, the median position weighted by site usage defines the start or end of the 5′ or 3′ UTR respectively, for example CDS 2 3′ UTR. SLASs and PASs are assigned to a CDS if there are no intervening sites with high usage. For example, the SLAS in the candidate 3′ UTR of CDS 1 (bold) is ignored when assigning the higher usage PAS to the next upstream CDS, CDS 1. This SLAS is not assigned to CDS 2 due to the higher usage PAS between it and the next downstream CDS, CDS 2. **B.** Illustrative examples of when the SLAS assigned to a CDS suggests (top) that an in-frame upstream methionine codon should be the CDS start codon or (bottom) that an in-frame downstream methionine codon should be the CDS start codon.

To handle the end of co-transcribed units, SLASs and PASs from either strand block assignment of PASs and SLASs respectively. Assignment behaviour can be tuned by a usage factor (`slapassign --min-usage-factor`): If the relative usage of any intervening SLAS/PAS site to the usage of the candidate PAS/SLAS site is at least the usage factor, then it will block the candidate site from being assigned to a CDS. For a usage factor of 0.5, then any intervening PAS/SLAS site with usage of at least half of the candidate SLAS/PAS site will block assignment. For a usage factor of zero, then all intervening PAS/SLAS sites will prevent assignment of sites, irrespective of how infrequently observed. For a very large usage factor, bigger than the maximum site usage, then intervening PAS/SLAS sites are all ignored, and all sites up to the next CDS will be assigned.

We tested the impact of `--min-slas-usage` and `--min-pas-usage` on SLAS and PAS assignment to CDSs to select appropriate default values. We want SLASs and PASs to be dominated by true detections and reasoned that this can be tested by proportion assigned to a CDS – true sites should be readily assigned to a CDS while false positive sites would not necessarily be in biologically expected locations (eg. within a CDS, PASs with an intervening high usage SLAS and vice versa). Testing a range of minimum usage values showed a marked increase in assigned SLASs and PASs (measured by both number of sites and sum usage of the sites) from 1 to 4, before broadly plateauing (Figure 4A), therefore we set a default of 4 for both.

**Figure 4.**
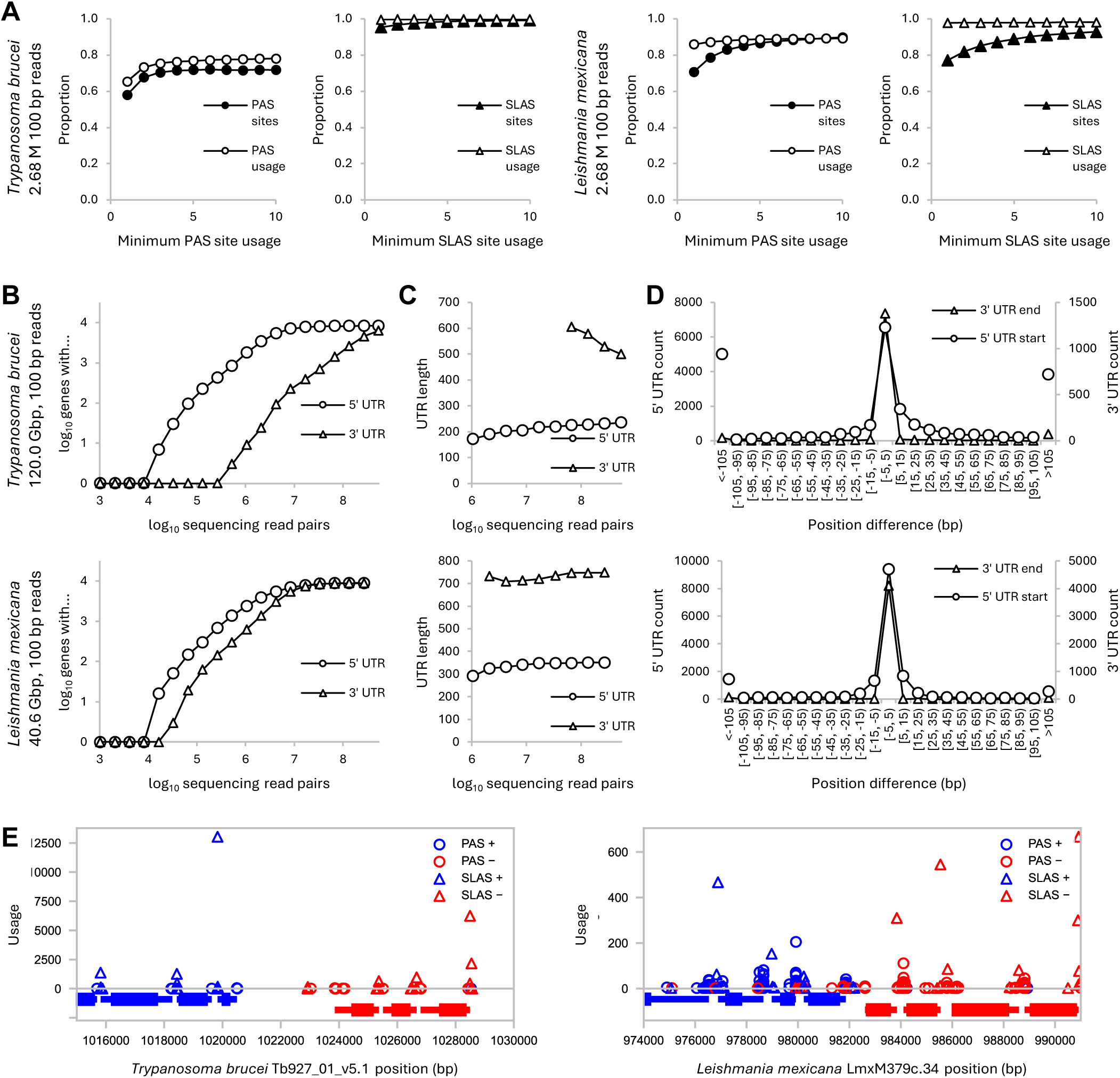
Benchmarking 5′ and 3′ UTR assignment rate **A.** Proportion of PASs and PAS usage assigned to a CDS for different minimum PAS usage and proportion of SLASs and SLAS usage assigned to a CDS for different minimum SLAS usage. Determined for (left) *T. brucei* and (bottom) *L. mexicana* using paired end 100 bp read sequencing data. **B.** The number of 5′ UTRs and 3′ UTRs assigned to a CDS for increasing numbers of read pairs. Detected using 6 bp and 9 bp match length for PAS and SLAS detection, as in Figure 2. Determined for (top) *T. brucei* and (bottom) *L. mexicana* using paired end 100 bp read sequencing data, for different numbers of randomly subsampled sequencing reads. **C.** The mean length of 5′ and 3′ UTRs, for different numbers of randomly subsampled sequencing reads, plotting data points where at 1/8 CDSs were assigned the 5′ or 3′ UTR respectively. **D.** Difference in position of detected and previously annotated start and end of 5′ UTRs and 3′ UTRs respectively, comparing *T. brucei* to TriTrypDB v68 (Amos et al., 2022) and *L. mexicana* to Fiebig et al., 2015. **E.** Example PAS, SLAS and UTR annotations for example genomic loci in *T. brucei* and *L. mexicana,* selecting a converging boundary between two transcription units in both species. Gene models are represented at the bottom of the plot, where the thick line represents the CDS and the thinner line represents the UTR, using the same strand colour code as PASs and SLASs.

We did not optimise `--min-usage-factor`, leaving it with the default value of 1.0. Unlike the minimum usage values, we could not determine an unbiased way to select an optimal value (increasing it will always increase number of sites assigned and average UTR length, and vice versa). It also has relatively little effect on the assignment of most SLASs and PASs to CDSs but may be useful for fine-tuning results. Therefore, it is provided as an option.

Number and incidence of usage of SLASs and PASs for each gene may be an informative data output for some types of mutant or life cycle stage comparison analyses, therefore we included slapstats which provides these statistics from GFF file generated by this pipeline.

### Performance of UTR definition based on spliced leader and polyadenylation sites using slaputrs

Next, we tested the combined performance of slapassign and slaputrs to 3′ and 5′ UTRs to protein coding gene models based on SLASs and PASs assigned to the CDS.

First, using the *T. brucei* and *L. mexicana* datasets, we asked how the number of genes with an annotated 5′ or 3′ UTR depends on read number (Figure 4B). As expected, this increases approaching a saturation point, which was approaching the number of CDSs annotated in the genomes. Detection of both *L. mexicana* UTRs and *T. brucei* 5′ UTRs saturated well under total reads available, while detection of *T. brucei* 3′ UTRs was underpowered, consistent with lower powered PAS detection.

For low numbers of assigned UTRs, mean length may be skewed due to preferred detection of highly abundant transcript SLASs/PASs. However, once approaching annotation of all UTRs (we used the threshold of at least one in eight CDSs having been assigned a UTR) any such bias should be reduced. We asked whether length of UTR detected was dependent on read number, as accurate annotation should not change with excess data. We observed such independence (Figure 4C).

We next tested re-annotation of UTRs in species with existing UTR mapping – looking for agreement in calling the UTRs (Figure 4D). The first test case was the *L. mexicana* dataset, which was previously used to identify SLASs and PASs using SLaP mapper (Fiebig et al., 2014), assigned them by proximity to CDSs (Fiebig et al., 2015) and then used to map UTRs based on the most frequently used SLAS and PAS (Beneke et al., 2022). Our `slapquant`, `slapassign`, `slaputrs` pipeline annotated 8902 5′ and 8825 3′ UTRs for 9182 CDSs. Of the 8629 5′ and 8485 3′ UTRs which both we and the previous study called, we asked which were identical or similar: 8082 (93.6%) 5′ UTRs were identical (except for a −2 bp offset due to convention about recording splice site coordinates). 3′ UTRs were more variable, but 4705 were within 5 bases (55.4%) and 7492 were within 100 bases (88.3%). There are several possible causes of these small differences: First, the RNA sequencing data is from the *L. mexicana* strain MNYC/BZ/62/M379 but the original identification of SLASs and PASs used the *L. mexicana* MHOM/GT/2001/U1103 reference genome. Second, original SLAS and PAS site identification used the SLaP mapper (Fiebig et al., 2014) approach, which differs from `slapquant`. Third, UTR calling in the original study used only the most frequently used SLAS and PAS rather than using any averaging, which `slaputrs` uses.

The second test case was annotating *T. brucei* (Figure 4D), the only species to have comprehensive UTR annotations available on TriTrypDB. For *T. brucei*, our `slapquant`, `slapassign`, `slaputrs` pipeline annotated 8343 5′ and 6539 3′ UTRs for 9660 CDSs, somewhat more 3′ UTRs than previously annotated. Of the 8276 5′ and 4415 3′ UTRs which both we assigned and TriTrypDB provides, we asked which were identical or similar: 7192 (86.9%) 5′ UTRs were identical. 3′ UTRs were more variable, but 1227 were within 5 bases (27.8%) and 2755 were within 100 bases (62.4%). As the origin of the reference UTR annotation is not well documented, this 3′ UTR discrepancy is harder to analyse. It may be genuine biological variability, for example derived from data from a different life stage, under different culture conditions, change in usage of PASs over time in culture, or properties of the methodology previously used.

Suspicious of this performance on *T. brucei*, we critically analysed analysis parameters. We tested a range of minimum PA match length and minimum PAS site usages with no obvious change to this measure – the 3′ UTRs detected always had comparable variability relative to the reference genome annotated UTRs. Similarly, gathering more input sequencing data and selecting different dataset types (poly-T vs. randomly primed for reverse transcription, data from our previous vs. other research group’s studies, data listed as strain-specific TREU927 vs. general *T. brucei* data, using multiple life cycle stage sequencing data) did not make a clear change. The accuracy of *L. mexicana* and consistent 3′ UTR discrepancy in *T. brucei,* combined with plausibility of UTRs assigned based on manual inspection (eg. Figure 4E), suggests that the reference genome 3′ UTR annotations may be incorrect for a subset of genes.

### CDS correction using SLAS and PAS sites

In CDS-first genome annotation, assigning the correct start codon for a CDS from genome sequence alone is challenging. There may be alternate methionine codons in-frame up and downstream of the selected start codon which could plausibly be the true start codon, particularly for proteins with high sequence divergence among species. Presence of a SLAS within the CDS indicates that the next downstream methionine codon is a more likely start codon. Similarly, presence of an in-frame methionine codon upstream of the called start codon and before a SLAP suggests that the upstream methionine codon is the start codon.

In `slaputrs` we implemented the `--fix-shorten-cds` and `--fix-lengthen-cds` options which catch these cases and adjust the CDS, and thus the 5′ UTR, accordingly. This adjustment is based on the position of the start of the 5′ UTR which would be assigned, based on the usage weighted median position of the SLAPs upstream of the stop codon which were assigned to the CDS (Figure 3B). Both lengthening and shortening select the start codon by minimising the length of the resulting 5′ UTR.

### Detecting unprocessed transcript using slapspan

RNA is not immediately trans-spliced and polyadenylated after transcription, and there is a window of time prior to these processes. These unprocessed and potentially nascent transcripts can be detected in RNA-seq datasets, even if sample preparation (eg. PA-selection) was not optimised for their detection. Alignment of reads from such transcripts to the genome can span SLAPs and PASs, as these sites have not yet been modified. Information about SLAS and PAS site usage can be readily extracted from the output of `slapquant`. Therefore we developed a tool, `slapspan`, which quantifies incidence of SLAS and PAS-spanning reads, using the strategy we previously described to analyse changes in co-transcribed variant surface glycoprotein (VSG) and VSG expression site associated genes in *T. brucei* (López-Escobar et al., 2022). Complexity arises as there may be multiple SLASs and/or PASs per gene, and alignment of a read to the genome may span one or multiple sites, and sites may be widely spaced such that a read is not long enough to span all SLAS or PAS associated with a gene. Therefore, two statistics are reported, a raw count and a weighted count. The former is the sum of the number of times a SLAS associated with the gene are spanned by a read, and similar for the PAS. The latter is the same sum but weighted by proportion of times that SLAS or PAS site was observed as used for that gene in the reference GFF.

Analysis of reads which may originate from nascent transcript allows insight into transcriptional and co-transcriptional control of transcript abundance. *T. brucei* mRNA decay kinetics suggests co-transcription degradation of some transcripts (Fadda et al., 2014). Furthermore, for expression of variant surface glycoprotein (VSG) genes and the co-transcribed expression site associated genes (ESAGS), specific regulators are being discovered: Regulation of transcript abundance from these specialised RNA Pol-I-transcribed expression sites, of which one is active at any one time, involves a transcriptional activator (ESB1) (López-Escobar et al., 2022) and ESAG transcript degrading factor (ESB2) (Lansink et al., 2025).

To test `slapspan` performance we re-analysed the effect of knockdown of ESB1 on PAS and SLAS-spanning reads, where we previously carried out a similar analysis albeit using a custom pipeline (López-Escobar et al., 2022). We annotated UTRs in the *T. brucei* Lister 427 genome using the control RNA-seq data from the study and the `slapquant`, `slapassign` and `slaputrs` pipeline, then measuring SLAP and PAS-spanning reads using `slapspan`. This reanalysis gave the same result as previously published, a decrease in PAS and SLAS-spanning reads for genes in the expression site (Figure 5A), consistent with ESB1 being required for active expression site transcription (López-Escobar et al., 2022). ESB2 is an endonuclease required for the lower level of relative to VSG from the active expression site, whose endonuclease activity and localisation to the expression site nuclear body suggests that it causes co-transcriptional degradation of some ESAG gene transcripts (Lansink et al., 2025). Using `slapspan` to analyse the published RNA-seq data from cells following ESB2 knockdown is consistent with this hypothesis, showing a decrease in PAS and SLAS-panning reads for ESAGs from the active expression site (Figure 5B). These results indicate `slapspan` is a useful addition to the toolkit for future related studies.

**Figure 5.**
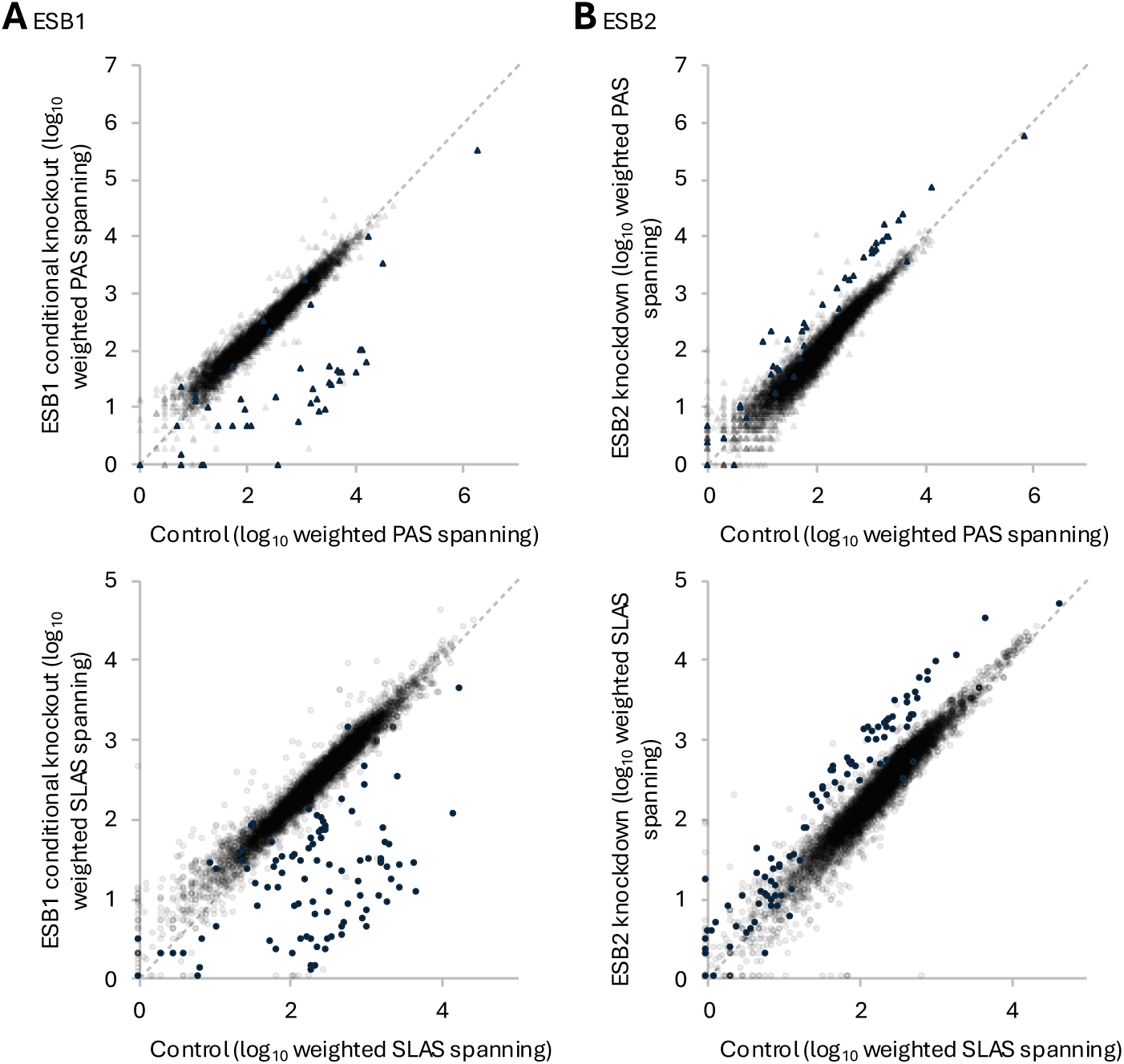
Detecting changes in PAS and SLAS-spanning reads in *T. brucei* mutants. Changes in PAS and SLAS-spanning reads in *T. brucei* Lister 427 parasites mutants, aligned to the 2018 genome resequence (Müller et al., 2018). **A.** Conditional knockout of ESB1, 48 h after induction of the phenotype. Data from López-Escobar et al., 2022. **B.** RNAi knockdown of ESB2, 24 h after induction of the phenotype. Data from (Lansink et al., 2025). For each, the top plot Is PAS usage-weighted number of PAS-spanning reads, and the bottom is SLAS usage-weighted number of SLAS-spanning reads. Each open data point represents a single transcript. Filled points are transcripts from a bloodstream form expression site (expression site associated genes, ESAGs, and expression site variant surface glycoprotein genes).

### Database-wide annotation of UTRs using the slapquant workflow

Having established the above method for UTR annotation, we aimed to provide UTR annotations for diverse species. We targeted every genome with CDS annotations in TriTrypDB version 68, 50 out of 88 genomes covering a range of human pathogenic and non-human pathogenic species and free-living relatives.

We set up a Snakemake (Mölder et al., 2025) workflow to find existing RNA sequencing data in the European Nucleotide Archive (ENA)/NCBI Sequencing Read Archive (SRA) where the recorded NCBI taxon ID matched the exact strain or, failing that, the species of each TriTrypDB genome. Aware that sequencing data may come from different life stages, or samples with perturbed expression due to reverse genetics mutants or chemical/drug treatment, we opted to carry out deep random data sampling. Up to 50 short read RNA-seq sequencing runs were selected at random. The workflow trims adapters from the reads, carries out a trial *de novo* transcript assembly using RNAbloom2 (Nip et al., 2020), from which the SL sequence is identified as we previously described (Wheeler, 2021b). When this failed, to avoid falling back on `slapquant` automatic SL sequence detection, we provided a manually specified sequence. Then, the workflow runs the `slapquant slapassign slaputrs` pipeline to annotate SLAPs, PASs and UTRs.

Of the 50 genomes, three failed due to technical issues, specifically due to the input GFF genome annotation file which contains CDSs not adhering to the GFF file format specification. For the remaining 47 genomes, there were a few problematic species, with near-complete failure to identify any UTRs in *T. brucei gambiense* DAL927, and near-complete failure to identify 3′ UTRs in *Paratrypanosoma confusum* CUL13, *Porcisia hertigi* MCOE/PA/1965/C119;LV43, *Endotrypanum monterogeii* LV88, *Leptomonas pyrrhocoris* H10. *T. brucei* TRUE927 was initially problematic, as it is a recently added NCBI taxon with limited strain-specific data indexed. However, we achieved high depth *T. brucei* TREU927 and *T. brucei gambiense* DAL927 UTR annotation by using more general data listed under *T. brucei*.

For our final analysis of the 47 genomes without technical problems, at least half of the CDSs were successfully assigned a 5′ UTR in 44 (93.6%) genomes, and a 3′ UTR in 31 (66.0%) genomes (Figure 6A). This is the first high coverage annotation of UTRs across the trypanosomatids.

**Figure 6.**
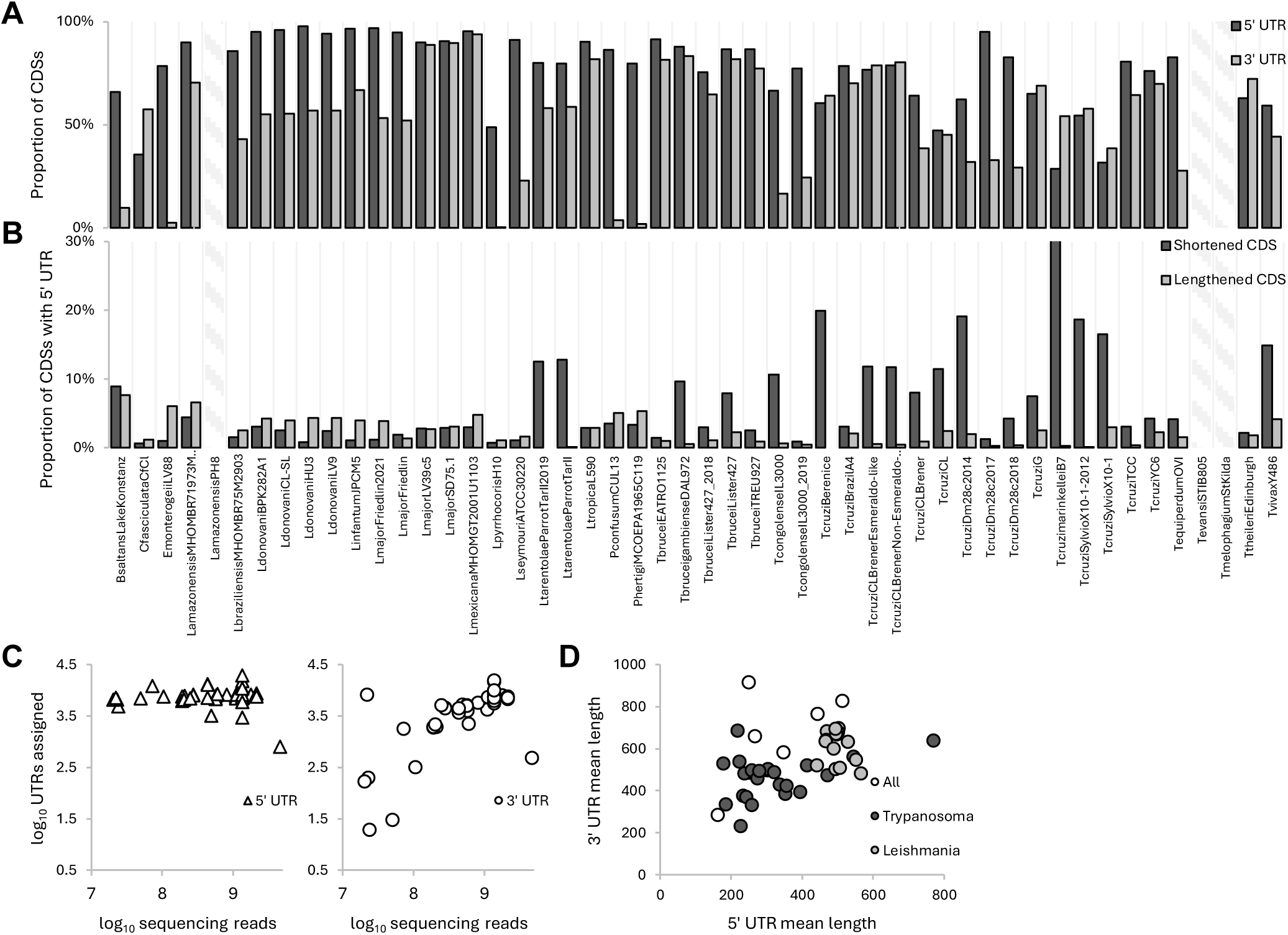
UTR annotation of trypanosomatid genomes using publicly available RNA sequencing data. **A.** Assignment of 5′ and 3′ UTRs to all 50 trypanosomatid genomes with CDS annotations available on TriTrypDB, using publicly available RNA sequencing data available from NCBI Sequencing Read Archive (SRA) or the European Nucleotide Archive (ENA). Bars represent the proportion of CDSs assigned a 5′ or 3′ UTR. Hatched columns indicate where UTR assignment was not possible due to issues with the input GFF files. **B.** Proportion of CDSs with SLAS(s) assigned enabling 5′ UTR assignment which were shortlisted for either start codon adjustment to lengthen the CDS based on an in-frame upstream start codon within the UTR, or CDS shortening to the first in-frame start codon downstream of the start of the assigned 5′ UTR. Hatched columns indicate where UTR assignment was not possible due to issues with the input GFF files. **C.** Correlation of performance in detection of 5′ and 3′ UTRs with input number of sequencing reads. Each data point represents one species from A. **D.** Correlation of 5′ and 3′ UTR length. Each data point represents mean 5′ and 3′ UTR length for a species, averaged across all CDSs with both UTRs assigned. Calculated without shortening or lengthening of the corresponding CDSs.

We used this genome set to test how often the assigned SLAS sites suggested that the CDS should either be shortened or lengthened. As expected, few potential corrections to CDSs were detected for most genomes, typically <5% of CDSs (Figure 6B). There was a tendency for CDS lengthening in *Leishmania* species and CDS shortening in *Trypanosoma* species. We speculate that this may be an annotation founder effect, where initial CDS calling in *Trypanosoma* and *Leishmania* had slightly different biases, which propagated as the original annotations were used for annotation training. Several genomes had a large proportion (>10%) of CDSs with a recommended shortening, suggesting systematic issues with the CDS annotation strategies, being too inclusive when discriminating CDS from UTR sequence.

This set of genomes provides a comprehensive test of the performance of `slapquant`, the dependency of UTR annotation on input read number on a high diversity of genomic and transcriptomic data (Figure 6C). The number of 5′ UTRs assigned had little correlation with number of input sequencing reads over the range tested (all >10^7^ reads). Conversely, the number of 3′ UTRs assigned had a broad positive correlation with read number (Figure 6C). Robust 5′ UTR annotation across diverse species can therefore be achieved with comparatively little data, while high depth 3′ UTR annotation requires more care when using short read sequencing data.

### Biologically relevant features of UTRs

Our comprehensive UTR annotation allows easy comparison of UTR properties across kinetoplastids. First, we asked whether different lineages have characteristic UTR lengths, considering the per genome average 5′ and 3′ UTR length (without CDS correction based on SLAS position). This showed a distinctive pattern where *Leishmania* species tend to have longer UTRs than *Trypanosoma* species, with slightly longer mean 3′ UTR length and much longer (almost double) mean 5′ UTR length (Figure 6D). We also noted no obvious difference between related lineages which have retained or lost the RNAi machinery, specifically *L. braziliensis* MHOM/BR/75/M2904 vs. the other *Leishmania* spp. in this set.

This also allows comparison of the rate of change in 5’ and 3’ UTR sequence over evolutionary time relative to the rate of change of the protein sequence encoded by the gene. Using *Leishmania major* as the reference species and *Leishmania infantum, Leishmania tarentolae* and *Crithidia fasciculata* as progressively more divergent species, we saw little correlation of UTR sequence conservation with protein sequence conservation (Figure 7). 5′ UTRs tended to be a little more similar between species than the 3′ UTRs, but with both reaching little more than the average sequence identity of two random DNA sequences when comparing at the *Leishmania* to *Crithidia* evolutionary distance. Rapid divergence of UTR sequence compared to protein sequence is not unexpected as important long linear motifs have not readily been found in UTRs, and more general sequence features instead seem important, like uracil-rich polypyrimidine tracts (Trenaman et al., 2024). Comprehensive UTR mapping enables future analyses for cross species analysis of important sequence features.

**Figure 7.**
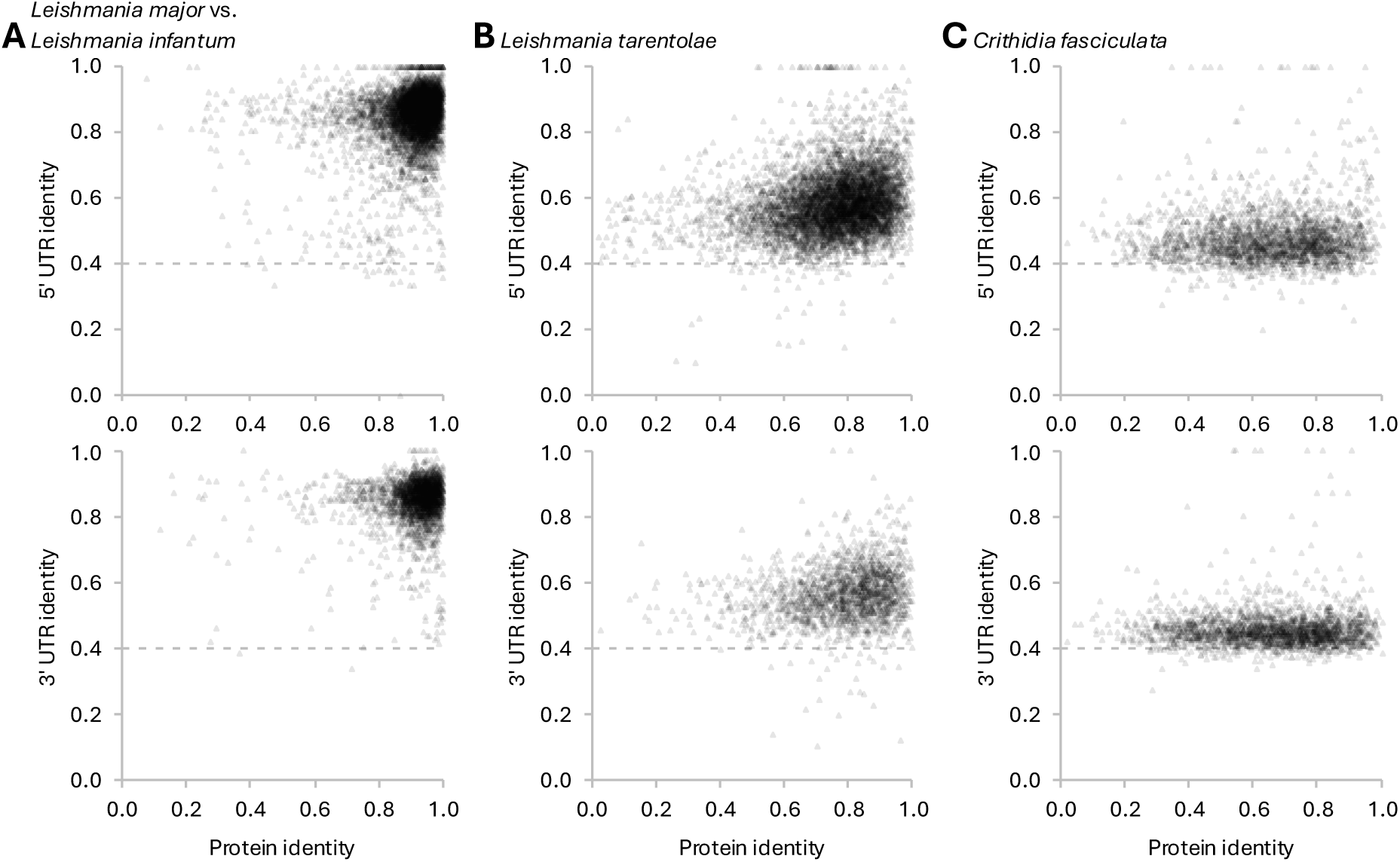
Sequence similarity of UTRs in comparison to protein coding sequence Correlation of protein sequence similarity with 5′ and 3′ UTR sequence similarity, comparing *Leishmania major* to a series of more divergent species: **A.** Comparison with *Leishmania infantum* (different *Leishmania* sub-genus). **B.** With *Leishmania tarentolae* (non-mammalian infective *Leishmania*). **C.** With *Crithidia fasciculata* (non-vertebrate infective *Leishmania* relative). Each data point represents the sequence identity (number of identical residues within the aligned portion, using Clustal Omega) of the protein coding sequence and nucleotide UTR sequences of a pair of transcripts, identified as orthologs by reciprocal best BLAST hit. The horizontal dashed line represents average sequence identity of two random DNA sequences.

## Discussion

Recognising a consistent gap in the annotation of trypanosomatid parasite genomes, we have developed, optimised, benchmarked and applied a practical tool for annotating SLAS, PAS, 3′ UTRs and 5′ UTRs from commonly available short read RNA sequencing data. Our approach follows similar approaches and ideas to previous tools, specifically SLaP mapper and UTRme (Fiebig et al., 2014; Radío et al., 2018), with two key technical adjustments. First, using clipping of aligned reads rather than filtering of reads by SL or PA sequence then alignment, to avoid the necessity of post alignment filtering or scoring. Second, carefully designed heuristics for UTR annotation from SLASs and PASs.

Perhaps more importantly, the Python-based `slapquant` tool is designed to be readily usable, only requiring AWK and BWA or BWA-MEM2 as dependencies, and be performant by avoiding writing of large intermediate files. This makes it straightforward and quick to run even on small workstations. It is sufficiently robust so that it can process large scale datasets of many genomes via data pipeline tools like Snakemake (Mölder et al., 2025) in reasonable time.

Having optimised settings on two test datasets, selecting one *Leishmania* and one *Trypanosoma* species to avoid over-fitting to any biological properties specific one of these human disease-relevant lineages, we demonstrate that slapquant can be used for SLAS, PAS and UTR annotation across almost all available trypanosomatid genomes currently available.

We hope that this will be adopted as a practical analysis step in trypanosomatid genome annotation, following annotation of CDSs using tools like AUGUSTUS (Stanke et al., 2004) or BRAKER (Brůna et al., 2021). In addition to informing the start of 5′ UTRs, SLAS mapping allows informed adjustment of CDSs. The start of the transcript is a biologically well-motivated means of determining start codon. As SLAS identification and thus 5′ UTR assignment using `slapquant` appears particularly reliable we suggest that this should be widely used to adjust predicted CDSs.

One complication with identifying SLASs is that the SL sequence must be known. We are aware of one dedicated SL sequencing finding tool SLFinder (Calvelo et al., 2020), and ourselves have developed a naïve tool specifically for the discoba lineage (which includes the trypanosomatids) (Wheeler, 2021a), which can overcome this hurdle. The automatic SL sequence identification which `slapquant` provides may also be sufficient but should be used only with caution.

Wide availability of UTRs provides many opportunities for future work. It provides the foundation for analysis of UTR contributions to transcript stability, life cycle regulation and translation efficiency, which go on to define the expression level of proteins encoded by trypanosomatid genomes. We anticipate applications in many studies, for example looking for RNA binding protein binding sites in UTRs involved in life cycle stage-specific expression, general transcript properties which modulate stability and thus overall expression level, and 5′ UTR features like short upstream ORFs which modulate translation efficiency.

The value of UTR annotation extends beyond these direct functional considerations. The ability to quantify transcripts from RNA-seq data by using full transcript, rather than just the CDS, enables more accurate abundance quantitation. This is particularly useful for discriminating between members of multi-copy gene families and tandem gene arrays, where CDSs may be highly conserved but UTR sequences have more gene-specific variation. Many single-cell RNA-seq technologies only sequence from the transcript 3′ end and accurately mapped 3′ UTRs enhance the ability to accurately interpret these data.

As with all analyses, high quality input data will ensure highest quality output. For genomes with little RNA sequencing data available, we observed a large variation in the ability to assign UTRs, particularly 3′ UTRs. We could not identify a clear correlation with sequencing run metadata (available in the SRA/ENA records), which suggests that precise RNA sample preparation and handling is impacting the usability of the resulting RNA sequencing data for calling UTRs. For UTR annotation, it is critical to ensure that best practices in preparing RNA samples are used, especially as detecting PASs is more challenging and 3′ to 5′ exonuclease activity is a major mRNA degradation pathway. Guided by previous carefully-designed analyses, such as de Freitas Nascimento et al., we recommend pelleting cells from culture in a warmed centrifuge, followed by immediate (without washing) pellet flash freezing or lysis, working with one sample at a time to minimise handling time.

Some of the variability in ability to identify SLAS, PAS and UTRs may, however, stem from true biological variability. The extent of true variation of SLAS and PAS site selection is not yet mapped, and may depend on precise culture conditions, life cycle stage, time in culture, strain or isolate-specific variation, etc. `slapquant` provides an effective tool for such future analyses.

There is also scope for future `slapquant` development, which we have not explored at this stage. For paired end sequencing data, it is expected that the SL sequence is only found at either the 5′ end of a read or its mate pair. Similarly, the reverse complement PA sequence is expected only to be found at the 3′ end of a read or its mate pair. We do not take this additional information into account and instead consider all provided reads in isolation. Specific paired end read handling, theoretically, has the potential to reduce noise in PAS identification twofold. Also, while SLAS sites tended to be very specific, PAS sites tended to have more variability. While genes tend to have a single cluster of preferred PASs this is not always the case, and occasionally there are two or more clear clusters. We do not attempt to call these alternate SLASs or PAS clusters as alternate transcripts. Finally, we have focused on high accuracy short read sequencing data, but the same SLAP, PAS and UTR assignment strategies could be used, with adaptation to handle lower sequencing accuracy, for long read and direct RNA sequencing technologies.

## Methods

### Data sources

Unless otherwise indicated, all genomes and genome annotations were fetched in FASTA and GFF format respectively from TriTrypDB (Amos et al., 2022), version 68. This includes the reference set of UTR annotations we used for *T. brucei* TRUE927. *Leishmania mexicana* MNYC/BZ/62/M379 expressing Cas9 and T7 RNA genome and genome annotations were previously described (Beneke et al., 2022), and fetched from the current version of the corresponding Zenodo data deposition (Beneke et al., 2025). This includes the reference set of UTR annotations we used for *L. mexicana*, the definition of which is described in Beneke et al., 2022: SLAPs and PASs were identified using SLaP mapper (Fiebig et al., 2014), UTRs for novel transcripts and with adjusted CDSs were defined in and taken from Fiebig et al., 2015, otherwise the most commonly used SLAP within 5 kb upstream and PAS 5 kb downstream were taken as the start of the 5′ and end of the 3′ UTR respectively.

Test RNA-seq datasets for general tests were manually selected. For *T. brucei,* the following SRA/ENA accessions from our study discovering ESB1 (López-Escobar et al., 2022), all from procyclic form *T. brucei* TREU927: SRR17052087 (Parental Cas9 and T7-expressing cell line); SRR17052090, SRR17052089, SRR17052088 (ESB1 deletion, identified as having no transcriptome-detectable defect); SRR17052096, SRR17052094, SRR19091051, SRR19091050, SRR19091048 (uninduced ESB1 overexpression cell line). For *L. mexicana,* the following SRA/ENA accessions from a previous transcriptomic analysis of wild-type *Leishmania mexicana* MNYC/BZ/62/M379 (Fiebig et al., 2015):

ERR789796, ERR789787, ERR789801, ERR789798, ERR789800 (promastigote); ERR789791, ERR789788, ERR789790, ERR789799, ERR789797 (axenic amastigote).

RNA-seq datasets for testing `slapquant` were manually selected from previous studies analysing ESB1 (López-Escobar et al., 2022) and ESB2 (Lansink et al., 2025), all from *T. brucei* Lister 427. ESB1 conditional knockout control (with induced exogenous copy): SRR17052110 SRR19091057 SRR19091056 SRR19091055. ESB1 conditional knockout phenotype induced (knockout after 48 h without induction of exogenous copy): SRR17052107 SRR19091054 SRR19091053 SRR19091052. ESB2 RNAi control (uninduced): ERR15084310 ERR15084311 ERR15084312. ESB2 RNAi phenotype induced (24 h induced RNAi): ERR15084313 ERR15084314 ERR15084315.

RNA-seq for annotation of TriTrypDB genomes was selected programmatically, based on sequencing run metadata available on ENA/SRA. We first manually selected a NCBI taxon IDs for each input genome, select an exact strain match when such a taxon entry was available and there was RNA-seq data in ENA/SRA listed under that taxon, otherwise using an exact species match. RNA-seq runs using paired end short read Illumina or BGI sequencing technologies matching each corresponding taxon ID were considered. A custom scoring scheme was used to prefer paired end data and avoid metagenomic or single cell sequencing data. Datasets which had very short reads (<40 bp average read length) or low read number (<100,000 reads) runs were not considered for selection. The remaining sequencing runs were selected, up to a maximum of 50 per genome.

### SLAP, PAS and UTR annotation

All new SLAP, PAS and UTR annotations described used the slapquant tools, using the settings specified in the text or figures.

The `slapquant` toolkit is available at https://github.com/Wheeler-Labab/slapquant. This is the suite of software tools which should be installed and used as explained in the corresponding documentation.

The scripts for testing and optimising the `slapquant` toolkit (presented in Figure 2 and Figure 4) is available at https://github.com/Wheeler-Lab/slapquant-tests. This is provided for reproducibility of the presented results, rather than for reuse.

The Snakemake (Mölder et al., 2025) workflow for fetching TriTrypDB genomes, selecting and fetching corresponding RNA-seq data, and for carrying out SLAP, PAS and UTR annotation is available at https://github.com/Wheeler-Lab/slapquant-annotations. This can be used to improve annotation of genomes as more RNA-seq data becomes available and can be extended for annotation of new genomes as they become available.

## Data availability

All software developed in this study, including slapquant, slapassign, slaputrs, slapspan, and slapidentify, is open-source and available on https://github.com/Wheeler-Lab/slapquant. The Snakemake workflow is available from https://github.com/Wheeler-Lab/slapquant-annotations.

The SLAS, PAS, UTR and corrected ORF annotations for the 47 trypanosomatid genomes generated during this study are available as GFF files through Zenodo (DOI 10.5281/zenodo.18759183). All raw RNA-sequencing data used for these annotations were obtained from public repositories (ENA/SRA); specific accession numbers for the datasets sampled for each genome are listed in **Supplementary Table S1**.

## Acknowledgements

This work was supported by the Wellcome Trust [Grant number 221944/20/Z].

We would like to thank Joana Faria (University of York) for constructive discussions concerning the ESB2 data.

## Copyright statement

For the purpose of open access, the author has applied a CC BY public copyright licence to any Author Accepted Manuscript version arising from this submission.

**Table S1. SRA accessions used for UTR annotation of genomes in TriTrypDB**

**Table S2. SLAS, PAS and UTR detection and assignment statistics for all TriTrypDB genomes with annotated CDSs.**

